# Neuropsychology of cognitive aging in rhesus monkeys

**DOI:** 10.1101/2023.05.30.542956

**Authors:** Mark G. Baxter, Mary T. Roberts, Jeffrey A. Roberts, Peter R. Rapp

## Abstract

Aged rhesus monkeys, like aged humans, show declines in cognitive function. We present cognitive test data from a large sample of male and female rhesus monkeys, 34 young (3.5-13.6 years) and 71 aged (19.9-32.5 years of age at the start of cognitive testing). Monkeys were tested on spatiotemporal working memory (delayed response), visual recognition memory (delayed nonmatching-to-sample), and stimulus-reward association learning (object discrimination), tasks with an extensive evidence base in nonhuman primate neuropsychology. On average, aged monkeys performed worse than young on all three tasks. Acquisition of delayed response and delayed nonmatching-to-sample was more variable in aged monkeys than in young.

Performance scores on delayed nonmatching-to-sample and object discrimination were associated with each other, but neither was associated with performance on delayed response. Sex and chronological age were not reliable predictors of individual differences in cognitive outcome among the aged monkeys. These data establish population norms for cognitive tests in young and aged rhesus monkeys in the largest sample reported to date. They also illustrate independence of cognitive aging in task domains dependent on the prefrontal cortex and medial temporal lobe.

(181 words)

## 1. Introduction

Macaque monkeys, especially rhesus monkeys, have been and continue to be a critical model for understanding the neurobiology of cognitive aging (Fletcher and Rapp, 2012; Gray and Barnes, 2019; Hara et al., 2012b; Morrison and Baxter, 2012). This is, in part, because of their highly differentiated prefrontal cortex relative to rodents, their sophisticated behavioral repertoire, and similarities between endocrine function and aging biology in macaques and humans.

Analyses of cognitive aging in humans have emphasized either dissociations between particular domains, such as item and source memory (Glisky et al., 1995), or central factors that drive functional decline across domains such as reduced processing speed with increased chronological age (Salthouse, 2010).

Against this background, the presence of substantial individual differences in cognitive aging is notable, with some older individuals performing well within the range of younger subjects while others show substantial impairment. This phenomenon is present in cognitive function in animals as well (Fletcher and Rapp, 2012; Gallagher and Rapp, 1997; Morrison and Baxter, 2012), suggesting that it reflects a biological aging process rather than the development of neuropathology unique to humans. This has led to the proposal that, like in humans, individual differences across behavioral domains in aged animals may be distinct, for example between sensorimotor tasks and spatial learning in aged rats (Burwell and Gallagher, 1993; Gage et al., 1984; Gallagher and Burwell, 1989) or between distinct cognitive domains (Barense et al., 2002; Beas et al., 2013; Gaynor et al., 2018; Johnson et al., 2017; LaSarge et al., 2007; Yoder et al., 2017). This has been less commonly investigated in aging in nonhuman primates (Bachevalier et al., 1991; Comrie et al., 2018; Frye et al., 2021; Gray et al., 2017; Herndon et al., 1997; Rapp and Amaral, 1989), in part because the number of monkeys usually available for characterization is relatively small. Whether age-related deficits tend to co-occur across functional domains versus showing independence has clear relevance for understanding biological mechanisms of cognitive aging, for example whether age-related cognitive decline is primarily attributable to a systemic factor like inflammation or metabolic dysfunction, or to region-specific alterations in neural function.

Here we report data from a large group of rhesus monkeys tested on multiple cognitive tasks using consistent apparatus and testing procedures, aggregating data collected over the course of a long-standing research program on neurocognitive aging in nonhuman primates. These tests include spatiotemporal working memory measured by the delayed response (DR) test, visual recognition memory measured by delayed nonmatching-to-sample (DNMS), and stimulus-reward association learning measured by the object discrimination (OD) test. This approach has several advantages. The sample size allows for a determination of reference norms on these tasks for research programs with geriatric nonhuman primates. These tasks are well-characterized neuropsychologically in terms of the effects of focal damage to many different brain areas in macaque monkeys, and indeed the neurology of memory in nonhuman primates has evolved around these and similar behavioral assessments (Murray and Baxter, 2006). DNMS is among the most thoroughly characterized tasks in behavioral neuroscience, at least with nonhuman primates. Broadly speaking, performance on the DR task depends on an intact prefrontal cortex (Bachevalier and Mishkin, 1986; Goldman et al., 1971), and performance on the DNMS and OD tasks require medial temporal lobe integrity (Meunier et al., 1993; Suzuki et al., 1993; Zola-Morgan et al., 1993).

## 2. Materials and Methods

### 2.1. Subjects

Data are presented from 105 rhesus monkeys (*Macaca mulatta*); 34 were “young”, 3.5-13.6 years old (mean, 7.6 years) at the beginning of delayed response testing, 21 female and 13 male, and 71 were “aged”, 19.9-32.5 years old (mean, 25.0 years), 41 female and 30 male. Although our primary analyses treat age as a binary “young” vs “aged” variable (Coleman et al., 2004), we note that the age range spanned the entire adult lifespan of rhesus monkeys, albeit with the omission of middle-aged animals (14-19 years).

Thirty-two of the 34 young monkeys (94%) and 28 of the 71 aged monkeys (39%) were born at the California National Primate Research Center. All monkeys included in this analysis were gonadally intact and none had received prior invasive or pharmacological manipulations expected to influence the cognitive or measures examined here. Most monkeys were singly housed at the time cognitive testing began, except for 5 young monkeys (1 male) and 6 aged monkeys (1 male). Formal food restriction schedules were not employed, but individual rations were adjusted to a level consistent with prompt responding in the test apparatus while maintaining healthy body weight and condition.

Testing took place in a manual test apparatus, identical to previous descriptions (O’Donnell et al., 1999; Rapp et al., 2003). A white noise generator was used throughout training to mask extraneous sound.

Acclimation and familiarization to the apparatus at the beginning of testing included offering the monkey food rewards in the test tray, as well as the opportunity to displace objects and plaques covering food wells in order to obtain reward. Behavioral data from some of the monkeys reported as part of this population have appeared in previous publications (e.g., Dumitriu et al., 2010; Hara et al., 2012a; Kyle et al., 2019; Long et al., 2020; Motley et al., 2018; O’Donnell et al., 1999; Shamy et al., 2011, 2006).

Testing took place in the following tasks in this order: delayed response, delayed nonmatching-to-sample, and object discrimination. All 105 monkeys began delayed response testing. 30 young and 61 aged monkeys continued to testing on delayed nonmatching to sample and all but 5 aged subjects continued to testing on object discrimination. 6 of the young monkeys had experience on a transitive inference task (Rapp et al., 1996) 2-3 years prior to delayed response testing; the others were naive to cognitive testing when delayed response testing began.

### 2.2. Delayed Response (DR) training and testing

DR training was conducted in phases, with and without delays. Training on the task rule took place with minimal memory demands, to ensure mastery of task structure before memory was challenged with increasing retention intervals. Trials were initiated by raising the opaque barrier of the apparatus, and the monkey watched through a Plexiglas screen while one of the lateral wells of the stimulus tray was baited with a food reward (e.g., raisin or peanut). The lateral wells were subsequently covered with identical plaques and, during the initial phase of training, the clear barrier was raised immediately to permit a response (0 s delay). After the monkey displaced one of the plaques, the opaque barrier was lowered to impose a 20 s intertrial interval (ITI). Daily test sessions consisted of 30 trials, with the left and right food wells baited equally often according to a pseudorandom sequence. Monkeys were trained until they achieved a criterion of 90% correct (9 errors or less in 9 consecutive blocks of 10 trials). Testing subsequently continued in an identical fashion except that a 1 s delay was imposed between the baiting and response phase of each trial, and continued until criterion performance was re-achieved. In the next phase the memory demands of the DR task were made progressively more challenging by introducing successive delays of 5, 10, 15, 30 and 60 s; testing was otherwise conducted as before (i.e. 30 trials/day, 20 s ITI). Monkeys were tested for a total of 90 trials (3 days of 30 trials per day) at each retention interval.

### 2.3. Delayed Nonmatching-to-Sample (DNMS) training and testing

Trials in this task consisted of two phases, initiated when the opaque barrier of the WGTA was raised to reveal an object covering the baited central well of the stimulus tray. After the reward was retrieved, the opaque screen was lowered, and the sample item positioned over one of the lateral wells. The other lateral well was baited and covered with a novel object. Like DR, training on the task rule took place with minimal memory demands before memory was challenged with increasing delay intervals. During training, a 10 s delay was imposed and recognition memory was tested by allowing monkeys to choose between the sample and the rewarded novel object. The discriminative stimuli were drawn from a pool of 800 objects according to a pre-determined sequence, ensuring that new pairs of objects were presented on every trial. Twenty trials per day were given using a 30 s ITI, counterbalancing the left/right position of the novel items. Subjects were tested until they reached a 90% correct criterion by committing no more than 10 errors in 5 consecutive sessions (100 trials). The memory demands of the DNMS task were then made progressively more challenging by introducing successively longer retention intervals of 15, 30, 60, 120 s (total = 100 trials each, 20/day), and 600 s (total = 50 trials, 5/day). Monkeys remained in the test chamber for all delays.

### 2.4. Object Discrimination (OD) testing

For each problem, two objects were positioned over the lateral wells of the apparatus, with their left/right locations varied pseudorandomly across trials. For each discrimination pair, one object was consistently associated with reward. A single “pre-trial” was run on the first session of each problem in which both objects were presented unbaited; the object the monkey did not pick on the “pre-trial” (which was not scored) was baited on the remainder of the trials for that problem. Test sessions consisted of 30 trials per day, using a 15 second intertrial interval. The first two test sessions were separated by 24 hours, and the third occurred after a 48-hour retention interval. Four discrimination problems were tested in this manner with the order of problems presented fixed across animals.

### 2.5. Measures of performance and data analysis

Trials to criterion, excluding the criterion run, were analyzed for acquisition of DR (at 0 s and 1 s delays) and DNMS (at 10 s delay) using one-way ANOVA with age group and sex as factors. Delay performance for DR and DNMS was analyzed using generalized linear mixed models on trial-level data, modeling binomial probabilities of correct vs incorrect responses on a trial as a function of delay, age group, and sex (fixed effects) and monkey identity (random effect). Performance on OD was analyzed based on average percent correct for each test session across the four discrimination problems, in a linear mixed model with test session, age group, and sex (fixed effects) and monkey identity (random effect). We did not use binomial regression for the OD data because the probability of a correct response would change within each test session as learning took place. Models included random intercepts for monkey. Statistical significance of main effects and interactions were determined by hierarchical likelihood ratio tests, building effects onto a null model with the random effect of monkey only. Correlations between task performance across monkeys were examined with both product-moment and rank correlations, using mean performance across delays (DR, DNMS) or problems (OD) as well as trials to criterion for DR and DNMS as summary measures. All analyses were performed in R 4.2.1 (R Core Team, 2022). The pattern of statistical results does not depend on using this more computationally intensive approach compared, for example, to repeated measures ANOVA, but is more flexible and accommodates the heterogeneity in error variance in performance measures based on an underlying binomial distribution (where variance is compressed at the upper end of the distribution of performance scores).

## 3. Results

### 3.1. Delayed Response (DR) training and testing

One aged female monkey (27.4 years old at the start of DR training) failed to reach criterion at the 0 s delay in under 1000 trials. Her testing was discontinued and this score was excluded from analysis of the 0 s training data. One aged female monkey was not trained at the 1 s delay, but proceeded from 0 s (criterion reached in 230 trials) to testing at the 5 s delay and continued with increasing delays as the other monkeys. Young monkeys attained criterion on the 0 s delay in a mean of 96 trials (range, 0-410) and aged monkeys in a mean of 106 trials (range 0-665), excluding the criterion run. Neither age nor sex, nor their interaction, accounted for a statistically significant proportion of variance in trials to criterion at 0 s, age F(1, 100) = 1.88, p = .17; sex F(1, 100) = 0.017, p = .90, age x sex interaction F(1, 100) = 2.44, p = .12. Young monkeys attained criterion on the 1 s delay in a mean of 96 trials (range, 0-614) and aged monkeys in a mean of 270 trials (range, 0-1990). The effect of age on trials to criterion at the 1 s delay was statistically significant, F(1, 99) = 10.76, p = .0014; the main effect of sex was not significant, F(1, 99) = 0.021, p = .89, nor was the age x sex interaction, F(1, 99) = 2.87, p = .093. Trends towards age x sex interactions in these measures, although not statistically significant, appeared to reflect slower acquisition of DR at both 0 and 1 s delays in aged female monkeys. Notably, the aged female monkeys were not older than the males on average. Not surprisingly, overall aged monkeys were more variable than young monkeys in trials to acquisition at 0 s and 1 s delays, F(33, 69) = 0.46, p = .016 and F(33, 68) = 0.13, p < .0001, respectively. Moreover, the aged female monkeys were more variable than the aged males on trials to criterion at both 0 s and 1 s delays, F(39, 29) = 5.03 and F(38, 29) = 9.19 respectively, ps < .0001. (This difference in variability did not compromise the ANOVA results for main effects of sex and age and their interactions; the pattern of results was identical when these analyses were done on rank-transformed data for which differences in variance between aged males and females were no longer statistically significant.) Summary statistics disaggregated by sex are presented in **Table 1**.

**Table 1.**
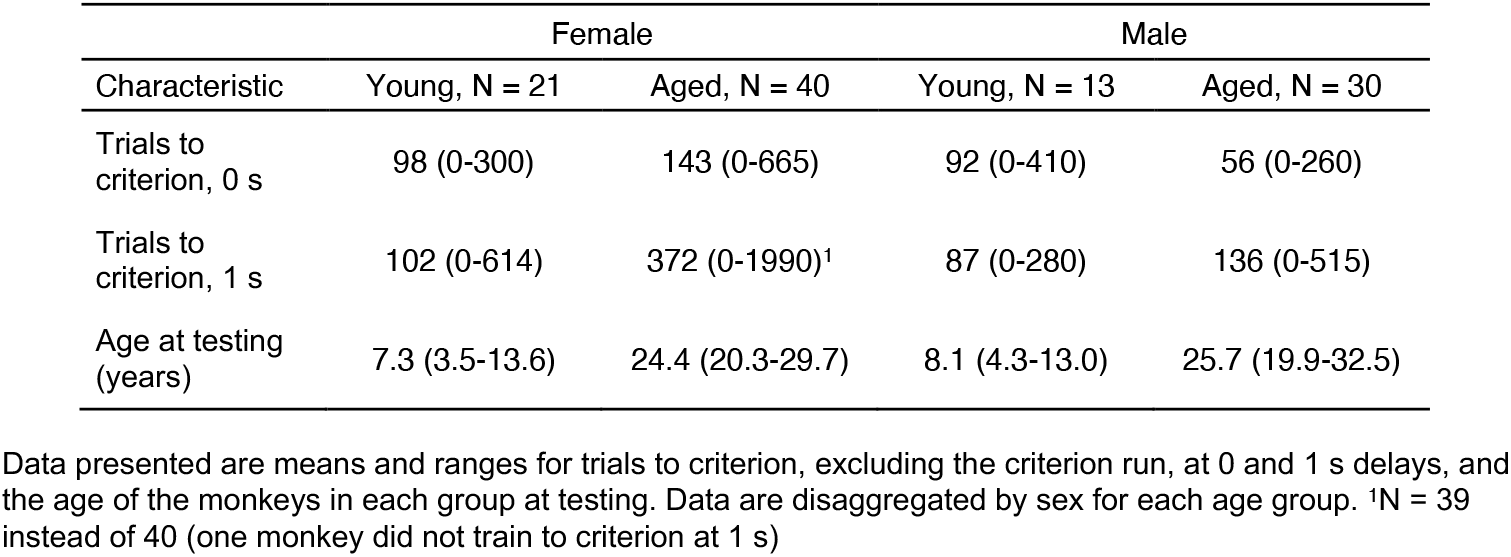
Delayed response training data.

| Characteristic | Female |  | Male |  |
| --- | --- | --- | --- | --- |
|  | Young, N = 21 | Aged, N = 40 | Young, N = 13 | Aged, N = 30 |
| Trials to criterion, 0 s | 98 (0-300) | 143 (0-665) | 92 (0-410) | 56 (0-260) |
| Trials to criterion, 1 s | 102 (0-614) | 372 (0-1990)<sup>1</sup> | 87 (0-280) | 136 (0-515) |
| Age at testing (years) | 7.3 (3.5-13.6) | 24.4 (20.3-29.7) | 8.1 (4.3-13.0) | 25.7 (19.9-32.5) |

Data presented are means and ranges for trials to criterion, excluding the criterion run, at 0 and 1 s delays, and the age of the monkeys in each group at testing. Data are disaggregated by sex for each age group. <sup>1</sup>N = 39 instead of 40 (one monkey did not train to criterion at 1 s)

Performance across 5-60 s delays was signficantly affected by delay, χ^2^(4) = 2124, p < .0001, age, χ^2^(1) = 25.6, p < .0001, and the interaction of delay and sex, χ^2^(4) = 23.1, p = .00012. The main effect of sex was not significant, χ^2^(1) = 3.50, p = .0612, nor were interactions of delay and age, χ^2^(4) = 5.59, p = .232, age and sex, χ^2^(1) = 0.07, p = .791, or the three-way interaction of age, sex, and delay, χ^2^(4) = 6.35, p = .174.

Analyses of estimated marginal means suggested that the interaction of sex and delay reflected poorer performance by female monkeys relative to males at the shorter delays, 5 s, z = -2.018, p = .0436; 10 s, z = - 3.008, p = .0026, but not at longer delays (15, 30, 60 s), |z| < 1.365, p > .17. The absence of interactions between age and sex, and of age, sex, and delay, suggests that although the detailed pattern of performance across delays between male and female monkeys may differ somewhat, age impacted the two sexes similarly. Delay performance is illustrated in **Figure 1.**

**Figure 1.**
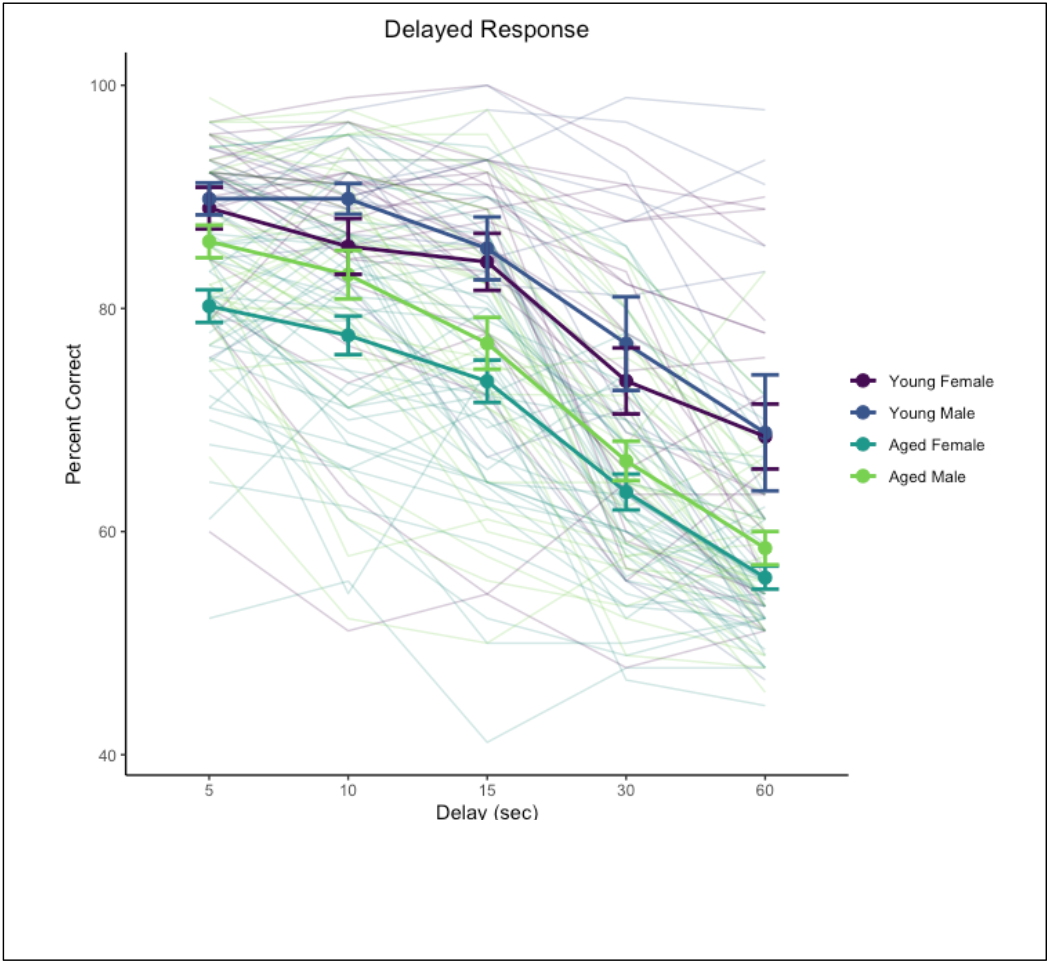
Delayed response performance across delay testing phase. Heavy lines indicate group means and standard errors, disaggregated by age and sex. Trace lines indicate performance of individual monkeys.

### 3.2. Delayed Nonmatching-to-Sample (DNMS) training and testing

Young monkeys acquired the DNMS task (at a 10 s delay) in a mean of 170 trials (range 20-440) and aged monkeys in a mean of 907 trials (range 12-1892). The main effect of age on trials to criterion was significant, F(1, 87) = 39.2, p < .0001, but effects of sex and the sex by age interaction were not significant, Fs < 1. Acquisition scores across the group of aged monkeys were more variable than young monkeys, F(29, 60) = 0.060, p < .0001. Summary statistics disaggregated by sex are presented in **Table 2**.

**Table 2.** Delayed nonmatching to sample training data

| Characteristic | Female |  | Male |  |
| --- | --- | --- | --- | --- |
|  | Young, N = 20 | Aged, N = 36 | Young, N = 10 | Aged, N = 25 |
| Trials to criterion | 166 (20-357) | 850 (12-1720) | 179 (20-440) | 988 (260-1892) |

Data presented are means and ranges for trials to criterion, excluding the criterion run, and are disaggregated by sex for each age group.

Performance across 15-600 s delays was significantly affected by delay, χ^2^(4) = 884, p < .0001 and age, χ^2^(1) = 31.8, p < .0001. The interaction of delay and age was not significant, χ^2^(4) = 2.44, p = .655. The main effect of sex was not significant, χ^2^(1) = 0.58, p = .45, nor were interactions sex with delay, age, or their interaction (χ^2^ < 1.17, p > .74). Delay performance is illustrated in **Figure 2**.

**Figure 2.**
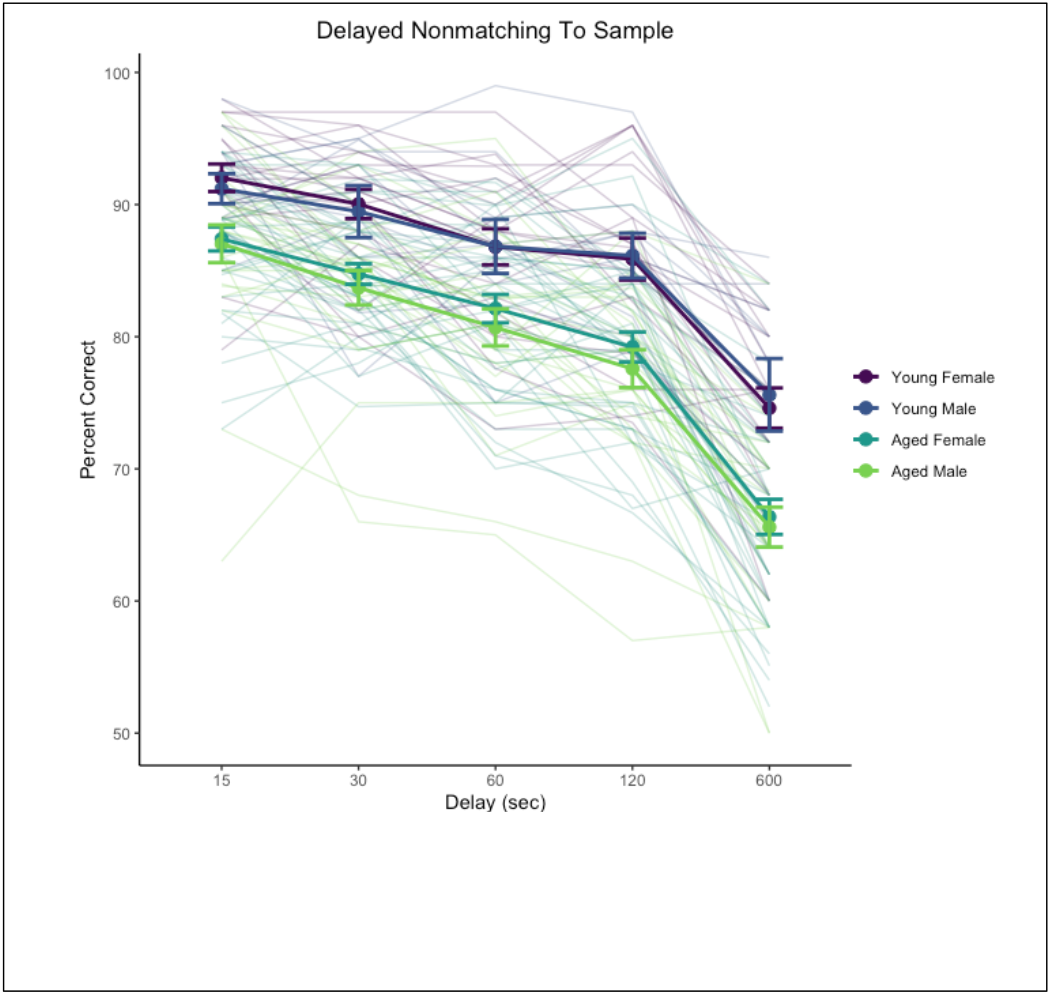
Delayed non-matching-to-sample performance across delay testing phase. Heavy lines indicate group means and standard errors, disaggregated by age and sex. Trace lines indicate performance of individual monkeys.

### 3.3 Object Discrimination (OD) testing

Monkeys learned new discrimination problems rapidly on the first day of training with each problem and improvement continued across subsequent days (**Table 3**). Overall performance scores (mean percent correct across delays for each monkey) did not show more variability in the group of aged monkeys compared to young monkeys, F(29, 55) = 0.58, p = .11.

**Table 3.** Object discrimination data

| Characteristic | Female |  | Male |  |
| --- | --- | --- | --- | --- |
|  | Young, N = 20 | Aged, N = 34 | Young, N = 10 | Aged, N = 22 |
| Day 1 percent correct | 85 (78-92) | 77 (56-90) | 79 (58-87) | 77 (60-91) |
| Day 2 percent correct | 95.6 (91.7-99.2) | 88.8 (71.7-99.2) | 93.5 (78.3-99.2) | 90.5 (72.5-98.3) |
| Day 4 percent correct | 98.2 (95.0-100.0) | 92.7 (77.5-100.0) | 96.2 (76.7-100.0) | 95.0 (85.8-100.0) |

Data presented are means and ranges for percent correct in each training session, averaged across the 4 problems presented. Data are disaggregated by sex for each age group.

Performance improved across test days, χ^2^(2) = 384, p < .0001, and was significantly affected by age, χ^2^(1) = 16.8, p < .0001. The interaction of day and sex was significant, χ^2^(2) = 9.21, p = .0099. The main effect of sex was not significant, χ^2^(1) = 0.027, p = .87, nor was the interaction of age and sex, χ^2^(1) = 3.35, p = .067, the interaction of day and age, χ^2^(2) = 4.56, p = .102, or the 3-way interaction of age, sex, and day, χ^2^(2) = 0.67, p = .72. Performance across days is illustrated in **Figure 3**.

**Figure 3.**
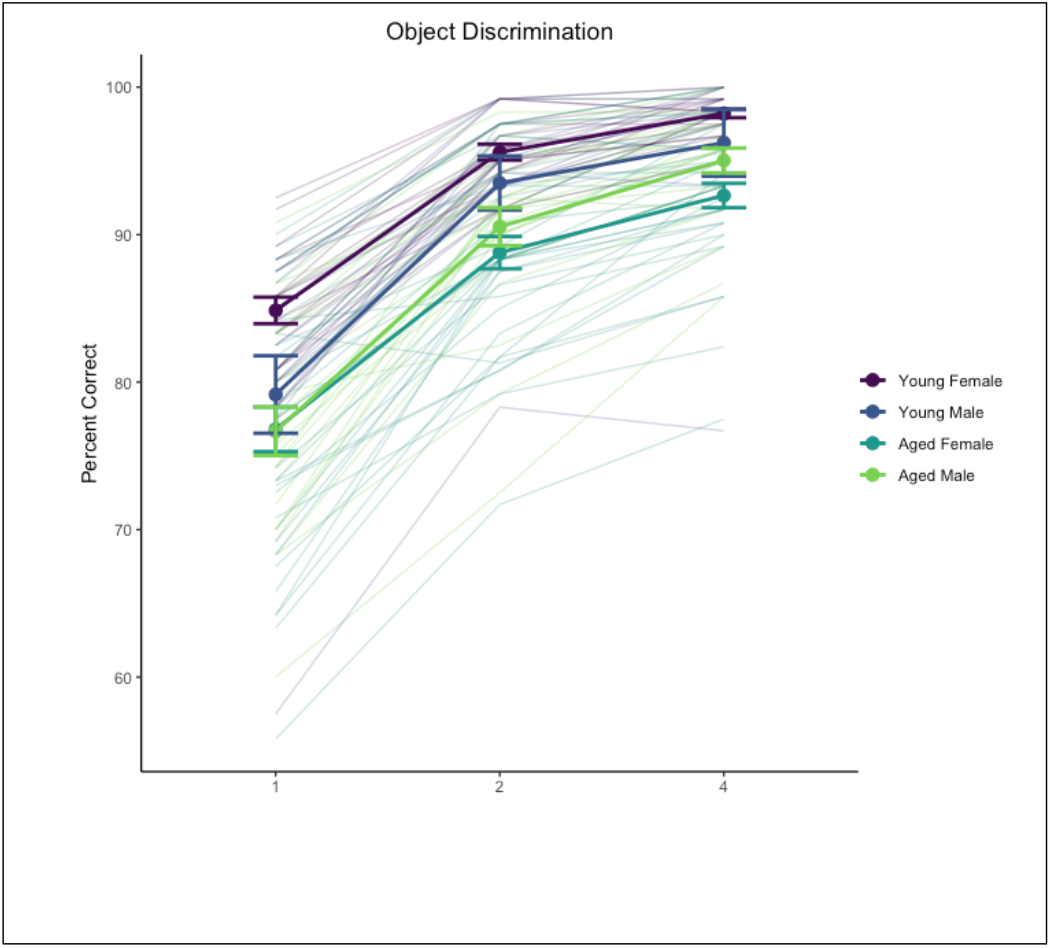
Object discrimination performance. Heavy lines indicate group means and standard errors, disaggregated by age and sex. Trace lines indicate performance of individual monkeys.

Analysis of the day by sex interaction with estimated marginal means suggested this was driven by an overall sex difference primarily on day 1 of testing with females outperforming males, t(125) = 2.05, p = .042, but not on days 2 or 4, |t(125)| < 0.137, p > .89. We also explored the age difference across test days separately for males and females, which suggested a more robust age effect on this task for females than males: effect of age on females t(125) > 3.29, p < .0013 for individual test days, whereas males did not show a robust age effect on any test day, t(125) < 1.295, p > .198. We interpret these differences with caution given the difference in sample size between males and females as well as the ceiling effect in discrimination learning where most monkeys performed nearly perfectly by the end of testing.

### 3.4. Relationship between performance across tasks

In general, performance on DR was unrelated to performance on DNMS and OD, but performance on DNMS and OD was positively associated, across the entire sample and among the aged monkeys alone. We considered six measures of performance: trials to criterion at 0 and 1 s delays in DR, average performance across delays in DR, trials to criterion in DNMS, average performance across delays in DNMS, and average performance across all days in OD. Spearman rank correlations between task measures are presented in **Table 4** for all monkeys and for aged monkeys alone.

**Table 4.** Spearman rank correlations between measures of task performance

| **All monkeys** | DR 0 s TTC | DR 1 s TTC | DR delays | DNMS TTC | DNMS delays | OD |
| --- | --- | --- | --- | --- | --- | --- |
| DR 0 s TTC | - | 0.06 | -0.11 | 0.04 | 0.03 | -0.07 |
| DR 1 s TTC | *0.55* | - | **-0.49** | 0.19 | -0.11 | **-0.26** |
| DR delays | *0.26* | *< .0001* | - | **-0.28** | 0.13 | **0.22** |
| DNMS TTC | *0.68* | *0.07* | *0.01* | - | **-0.61** | **-0.58** |
| DNMS delays | *0.77* | *0.28* | *0.22* | *< .0001* | - | **0.56** |
| OD | *0.51* | *0.02* | *0.04* | *< .0001* | *< .0001* | - |
| **Aged only** |  |  |  |  |  |  |
| DR 0 s TTC | - | 0.1 | -0.13 | 0.1 | 0.05 | -0.11 |
| DR 1 s TTC | *0.42* | - | **-0.49** | -0.05 | -0.02 | -0.1 |
| DR delays | *0.29* | *< .0001* | - | -0.01 | -0.02 | 0.06 |
| DNMS TTC | *0.43* | *0.68* | *0.92* | - | **-0.43** | **-0.37** |
| DNMS delays | *0.69* | *0.87* | *0.86* | *0.0005* | - | **0.4** |
| OD | *0.41* | *0.46* | *0.69* | *0.0053* | *0.0024* | - |

Correlations are presented for all monkeys and for aged monkeys only. Correlation coefficients are given above the diagonal, p-values (*italicized*) are below the diagonal (uncorrected for multiple comparisons; correlation coefficients $p < .05$ are in **bold**). TTC = trials to criterion. N = 86-105 (all monkeys) or 56-71 (aged only) per correlation.

Within DR, trials to criterion at 1 s correlated strongly and negatively with delay performance, suggesting that monkeys that found the 1 s delay challenging (higher trials to criterion) performed worse across delays (lower percent correct scores) even though they eventually achieved criterion performance at 1 s. Within DNMS and OD, trials to criterion on DNMS correlated strongly and negatively with delay performance on DNMS and percent correct on OD, which correlated positively with each other, again suggesting that monkeys that found DNMS acquisition challenging also showed poorer memory across longer delays in DNMS as well as worse discrimination learning performance. DR trials to criterion at 0 s did not correlate with any other measures of performance, suggesting this measure reflects some distinct factor (e.g., procedural learning capacity, neophobia or anxiety in acclimating to testing protocols). In the entire group of monkeys there were also moderate correlations between some DR measures and DNMS/OD measures, but these were not present in the aged monkeys alone, suggesting they might reflect a common main effect of age on these measures rather than an underlying relationship among them.

We considered whether classifying aged monkeys by performance on one task resulted in groups that were differentiated on other tasks (“impaired” / “unimpaired”) (Baxter and Gallagher, 1996; Gallagher and Burwell, 1989). We divided aged monkeys into “aged-unimpaired” and “aged-impaired” groups based on

DNMS performance, using the lowest average delay score for any monkey (78.0%) as the cutoff for impairment. The resulting aged-unimpaired and aged-impaired groups were significantly different from one another (z = 6.90, p < .0001) and both significantly differed from the group of young monkeys (mean DNMS delay performances young, 85.9%; aged-unimpaired, 82.5%; aged-impaired, 74.7%; z > 4.06, p ≤ .0001). We then reanalyzed the DR data using the same generalized linear mixed model structure as the primary analysis, but with 3 age groups (young, aged-unimpaired, aged-impaired) instead of 2. In this analysis, both aged groups were different from the young group, difference in estimated marginal means z = 3.00, 4.62 (ps < .008) for aged-unimpaired and aged-impaired respectively, but the aged-unimpaired and aged-impaired groups did not differ from one another, z = -1.12, p = .50 (indeed, the aged-impaired group numerically was better than the aged-unimpaired group). The reverse approach, splitting DR performance into impaired/unimpaired based on the minimum score in the young group, is problematic because DR performance is more variable in the young monkeys (worst delay performance across in a young monkey = 52.9%) so instead we used a median split of the aged monkeys to divide them into “above median” and “below median” groups as proxies for “unimpaired” and “impaired”. The resulting aged-above median DR and aged-below median DR were significantly different from one another (z = 6.61, p < .0001). The aged-below median DR group significantly differed from the group of young monkeys (z = 8.65, p < .0001) but the aged-above median DR did not (z = 2.10, p = .09); mean DR delay performances young, 80.9%; aged-unimpaired, 78.2%; aged-impaired, 65.5%). Applying this division of the aged monkeys based on DR to DNMS performance, both groups of aged monkeys were significantly different from young monkeys (z > 5.18, ps < .0001) but did not differ from each other (z = -0.43, p = .90).

Object discrimination performance shows a similar pattern to DNMS when split based on DNMS performance versus DR. These relationships are illustrated in **Figure 4**, suggesting that DR and DNMS/OD performance tap largely independent features of cognitive aging in monkeys.

**Figure 4.**
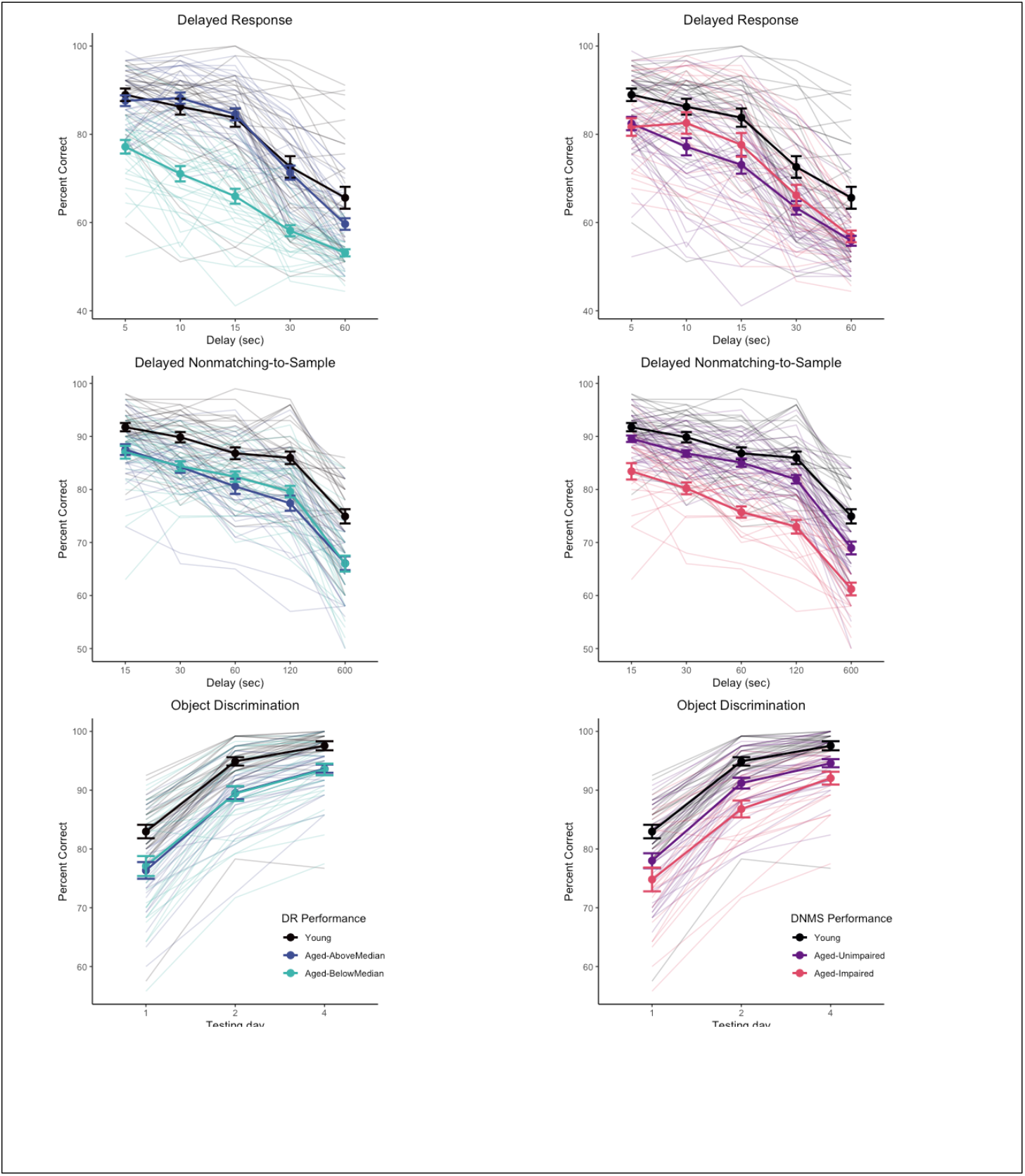
Task performance separated by “unimpaired” / “impaired” subgroups of aged monkeys based on DR (left column) or DNMS (right column) performance. Subgroups based on DR do not differentiate performance of aged monkeys on DNMS or OD, whereas subgroups based on DNMS differentiate performance of aged monkeys on OD but not DR. Heavy lines indicate group means and standard errors. Trace lines indicate performance of individual monkeys.

### 3.5 Does using age as a continuous variable make a difference?

The categorization of monkeys into “young” and “aged” is somewhat arbitrary, given that there is a wide age range in both groups, although there are no data in this study on monkeys between ∼13.5 and 19 years of age. We considered whether for primary analyses of delay performance on DR and DNMS, whether considering chronological age within “young” and “aged” groups would provide any more information than a categorical age variable alone. There was little relationship between chronological age and performance within the young and aged groups for either DR or DNMS, |r|s < 0.164, ps > .21. These findings underscore that while aging is a significant risk for cognitive impairment, age per se is a poor predictor of individual differences in the cognitive outcome of growing older.

### 3.6 Is there any evidence for a trend across time?

Because data on these monkeys were accumulated across a long period of time, we considered whether performance might vary as a function of the birth year of the monkeys, perhaps based on improvements in provision of environmental enrichment and social housing over time. Within the group of aged monkeys, birth year was not significantly associated with either average performance across DR delays (r^2^ = 0.02, p = .24) or DNMS delays (r^2^ = 0.005, p = .57).

## 4. Discussion

Three main conclusions emerge from considering the pattern of performance on these cognitive tasks in young and aged rhesus monkeys. First, age-related deficits in tasks broadly dependent on the prefrontal cortex versus the medial temporal lobe are independent of one another. Second, age-related deficits on tasks with different cognitive demands that share a dependence on the medial temporal lobe are correlated with each other. Third, there is little evidence for substantial sex differences in the pattern of age effects on this battery of cognitive tasks.

These data are, to our knowledge, the largest sample of cognitive data from young and aged rhesus monkeys reported to date. Previous investigations have either reported data on relatively small numbers of monkeys or have focused on global patterns of change across the lifespan (Herndon et al., 1997). The sample size and consistent testing of the same monkeys on a battery of cognitive tasks allows a determination of whether different behavioral tasks, which are all affected by aging, have coordinated or independent patterns of age effects (Bachevalier et al., 1991). A strength of this study is the use of behavioral tasks that are well-characterized in terms of the effects of focal lesions of different cortical and subcortical areas, and in the case of DR and DNMS, include training to criterion with minimal cognitive demands before challenging memory with increasing delays. This allows for clearer interpretation of the pattern of impairment because the ability to perform the task with minimal memory demands, even if more extensive training is required, indicates mastery of the task rules and intact sensory, motor, and motivational functions required to support performance. Our findings indicate independent patterns of age effects in different cognitive domains and, as such, call into question the approach of deriving a unitary composite score of cognitive performance to summarize cognitive ability in aged monkeys.

Monkeys whose cognitive data are included in this study have formed the basis for multiple investigations into the neurobiology of age-related cognitive decline (Dumitriu et al., 2010; Hara et al., 2012a; Long et al., 2020; Motley et al., 2018; Shamy et al., 2011, 2006). These investigations have identified patterns of change in synaptic parameters which differ between the prefrontal cortex and temporal lobe, against a background of preserved neuronal cell number. The independence of behavioral performance in tasks dependent on these two different cortical regions is consistent with differences in “synaptic strategy” underlying the neural computations in these different areas (Morrison and Baxter, 2012) and supports independent biological effects of aging on these different synaptic substrates. The absence of a pattern of dense cognitive impairment in aged monkeys, where all tasks are impaired in coordination, combined with evidence of preserved cortical neuron number argues against the presence of a naturally occurring dementia-like condition in aged rhesus monkeys (Beckman et al., 2019; Walker and Jucker, 2017).

There were subtle sex differences in performance across the tasks but age impacted performance similarly in both male and female monkeys. Female monkeys showed greater variability in acquisition of DR, likely related to variation in ovarian hormone status (Rapp et al., 2003). These monkeys were all gonadally intact and we did not attempt to determine menopausal status in females across the entire sample, although these data were available for a subset of monkeys that had participated in studies specifically aimed at determining the relationship between menopause and cognition (Roberts et al., 1997). The similarity in agerelated cognitive impairments on these tasks between male and female monkeys is reminiscent of the absence of sex differences across a battery of cognitive tasks in young make and female monkeys (Nagy et al., 2017).

This overall program of research was aimed at determining neurobiological substrates of impairments in “aged” monkeys compared to “young” monkeys rather than tracking patterns of behavioral and neurobiological changes across the entire lifespan. As such, the available data excludes middle-aged monkeys (roughly 14-19 years of age). Within the groups of young and aged monkeys, there was no relationship between chronological age at testing and performance, despite robust age differences between the two groups. Said another way, knowing whether a monkey is “young” or “aged” captures substantial variance in performance on these behavioral tasks, but it is not a reliable proxy for individual differences in cognitive outcome at older ages. Substantial individual differences were present within the group of aged monkeys, indicating that advanced chronological age is not synonymous with cognitive impairment. This implies that middle age in rhesus monkeys is a time of substantial cognitive and biological changes resulting in cognitive impairments in old age in some individuals. These changes are largely uncharacterized and represent an opportunity to understand the basis of age-related cognitive resilience versus impairment (Stern et al., 2023). Ultimately, longitudinal analysis will be critical for mapping life course trajectories of successful neurocognitive aging.

## Acknowledgments

We thank S. Fong, D. Kent, H. McKay, L. Novik, T. Ojakongas, C. Stanko for outstanding technical assistance throughout these studies, and Dr. C. A. Barnes for her contributions to this program of work.

This project was supported by NIH R01AG010606, R01AG003376, and in part by the National Institute on Aging Intramural Research Program.

